## Supplemental Data 1 for "ARHGAP18-ezrin functions as an autoregulatory module for RhoA in the assembly of distinct actin-based structures"

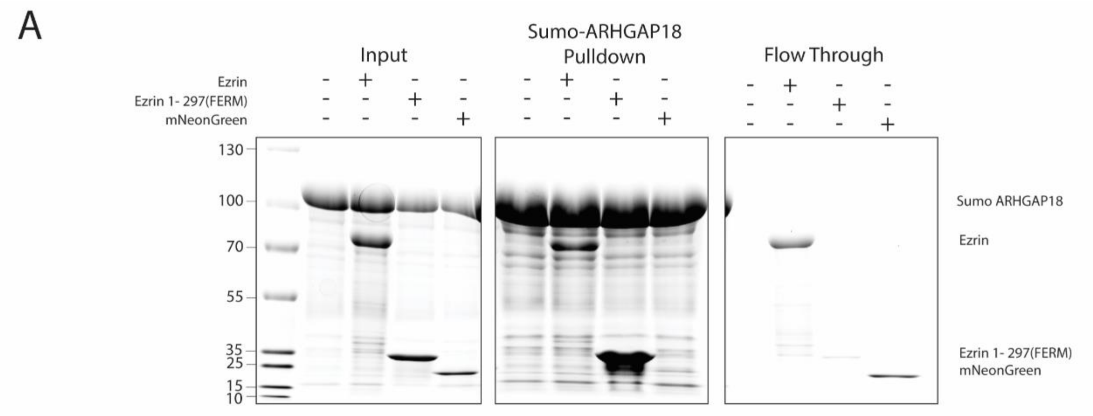


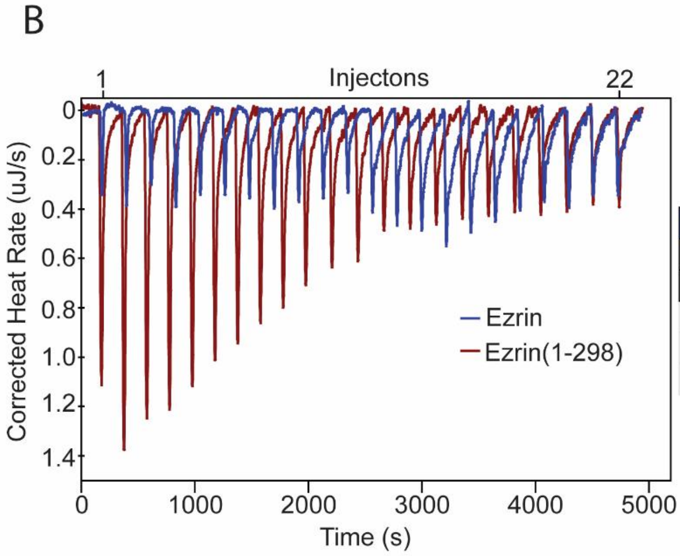


| Model | Variable | Mean Value (n=3) | SEM (n=3) |
|---|---|---|---|
| Blank (constant) | Blank (µJ) | -13.25 | 2.74 |
| Independent | Kd(M) | 19.29 | 3.53 |
| | n | 0.58 | 0.19 |
| | H(kJ/mol) | -45.51 | 21.93 |
| | S(J/mol*K) | -110.60 | 33.02 |

**Figure S1 ARHGAP18 affinity for ezrin using purified proteins.** A) Coomassie stained gels from assay testing the binding of Full length ezrin, ezrin-FERM (1-297) and mNeonGreen (negative control) to a resin saturated with SUMO-ARHGAP18. From left to right: Input proteins, pulled down proteins, flow through proteins. Both FL-ezrin and mNeon green are observed in the flow through while FERM is only found bound the column. FL-ezrin shows weaker affinity for the column than FERM whereas mNeonGreen did not bind the SUMO-ARHGAP18 column. B) Representative injections (above) and plotting of area under each injection peak with fit (below). FL-WT ezrin did not show measurable binding to ARHGAP18 while FERM produced measurable binding curves. Reported Kd is (mean ± SEM) from replicates of individual ITC experiments.

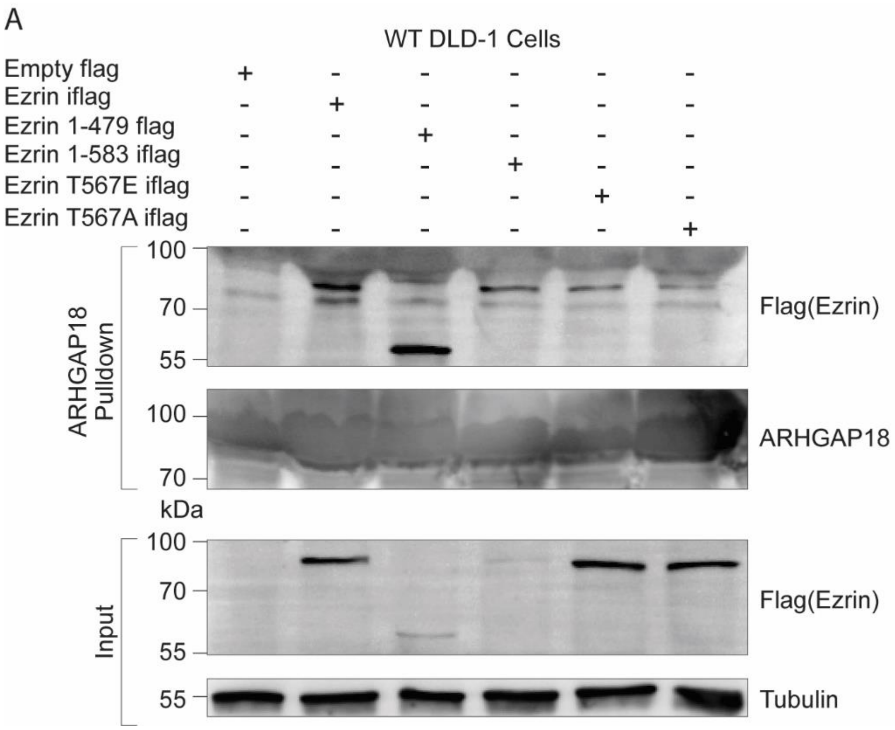


**Figure S2 ARHGAP18 binds active, open ezrin through the FERM domain in DLD-1 cells.** Western blot replication of experiment in Figure 1C except done in DLD-1 human colorectal cells instead of placental Jeg3 cells. Expression of constitutively active forms of ezrin and FERM is detrimental to many cell types leading to lower expression. Even with much reduced expression, comparison of input to pulldown shows a preferential binding of FERM to ARHGAP18 compared to WT ezrin.

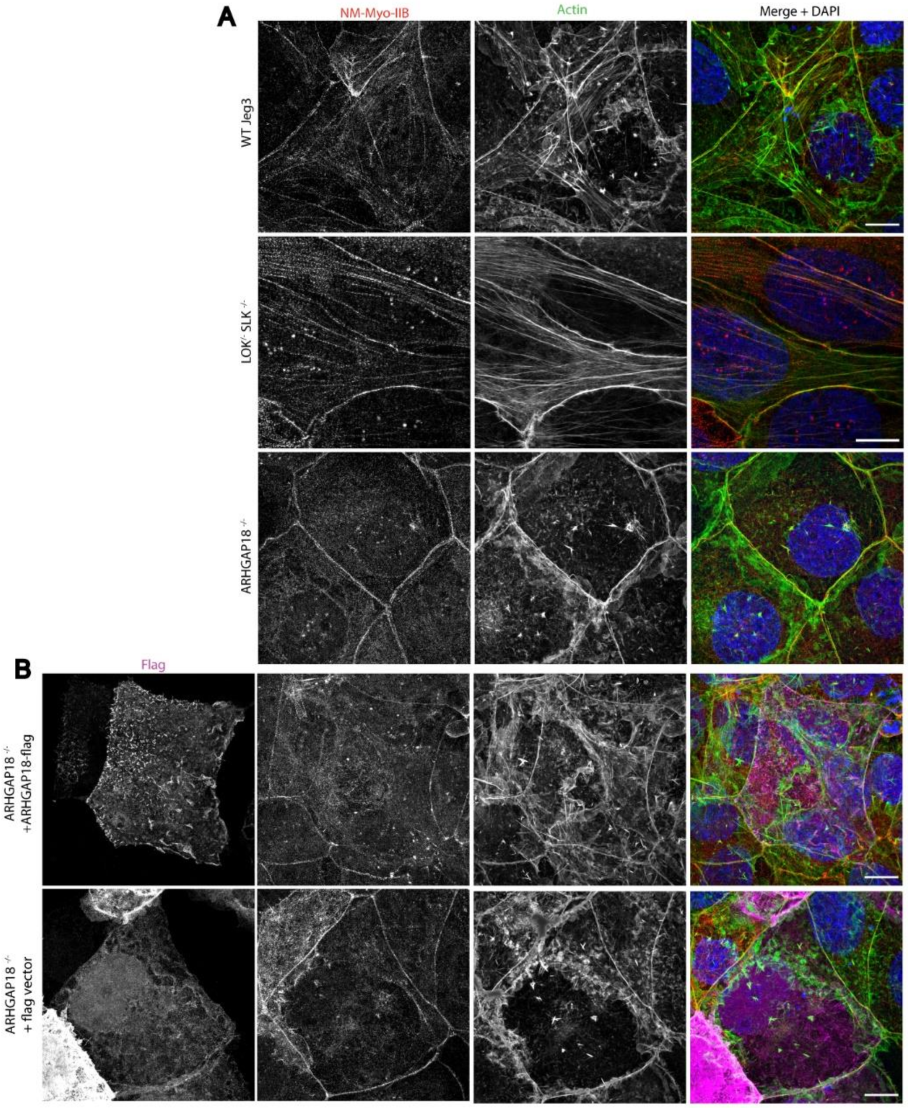


**Figure S3 Comparison of non-muscle Myosin IIb, and actin in WT Jeg3, LOK$^{-/-}$SLK$^{-/-}$ ARHGAP18$^{-/-}$, and ARHGAP18 rescue with empty vector control.** Representative maximum Z-projections of immunofluorescence SIM images of fixes cells. WT and ARHGAP$^{-/-}$ images are the same as main figure 5A shown here to present direct comparison to rescue, control and LOK/SLK knockout conditions. Large contractile fibers at the apical surface are increased in LOK$^{-/-}$ SLK$^{-/-}$ while reduced in the ARHGAP18$^{-/-}$ compared to WT Jeg3. Rescue of ARHGAP18$^{-/-}$ with a flag-tagged ARHGAP18 but not empty vector restores actin back to near WT conditions most clearly seen in the actin channel. Scale bars 10μm.

Movie 1: Time-lapse movie of live WT Jeg3 vs. ARHGAP18$^{-/-}$ cells transiently expressing GFP-EBP50 which targets to microvilli. Each frame taken at 1-minute intervals. ARHGAP18-/- cells have more but smaller microvilli, which turn over more rapidly than in the WT condition. Still frames of selected individual microvilli shown in Figure 3A. Scale bars 5µm.

Movie 2: Time-lapse movie of live WT Jeg3 vs. ARHGAP18$^{-/-}$ cells transiently expressing GFP-EBP50 which targets to microvilli. Each frame taken at 1-minute intervals. ARHGAP18-/- cells have more but smaller microvilli, which turn over more rapidly than in the WT condition. Still frames of selected individual microvilli shown in Figure 3A. Scale bars 5µm.

Figure 1-Source Data 1: Full Western blot images from all blots shown in Figure 1.

Figure 2-Source Data 1: Full Western blot images from all blots shown in Figure 2.

Figure 4-Source Data 1: Full Western blot images from all blots shown in Figure 4.
